# LANTERN: Leveraging Large Language Models and Transformers for Enhanced Molecular Interactions

**DOI:** 10.1101/2025.02.10.637522

**Authors:** Ha Cong Nga, Phuc Pham, Truong Son Hy

## Abstract

Understanding molecular interactions such as Drug-Target Interaction (DTI), Protein-Protein Interaction (PPI), and Drug-Drug Interaction (DDI) is critical for advancing drug discovery and systems biology. However, existing methods often struggle with scalability due to the vast chemical and biological space and suffer from limited accuracy when capturing intricate biochemical relationships. To address these challenges, we introduce **LANTERN** (Leveraging Large **LAN**guage Models and **T**ransformers for **E**nhanced molecula**R** interactio**N**s), a novel deep learning framework that integrates Large Language Models (LLMs) with Transformer-based architectures to model molecular interactions more effectively. LANTERN generates high-quality, context-aware embeddings for drug and protein sequences, enabling richer feature representations and improving predictive accuracy. By leveraging a Transformer-based fusion mechanism, our framework enhances scalability by efficiently integrating diverse interaction data while maintaining computational feasibility. Experimental results demonstrate that LANTERN achieves state-of-the-art performance on multiple DTI and DDI benchmarks, significantly outperforming traditional deep learning approaches. Additionally, LANTERN exhibits competitive performance on challenging PPI tasks, underscoring its versatility across diverse molecular interaction domains. The proposed framework offers a robust and adaptable solution for modeling molecular interactions, efficiently handling a diverse range of molecular entities without the need for 3D structural data and making it a promising framework for foundation models in molecular interaction. Our findings highlight the transformative potential of combining LLM-based embeddings with Transformer architectures, setting a new standard for molecular interaction prediction. The source code and relevant documentation are available at: https://github.com/HySonLab/LANTERN.

## 1 Introduction

Deciphering molecular interactions—including Drug-Target Interactions (DTI), Protein-Protein Interactions (PPI), and Drug-Drug Interactions (DDI)—is crucial for advancing drug discovery Sachdev & Gupta (2019); Liao et al. (2025), therapeutic innovation Grizzle et al. (2019); Luo et al. (2024), systems biology Hu et al. (2021); Meng et al. (2021), protein design Tran & Hy (2024); Nguyen et al. (2024), and protein-binding ligand generation Khang Ngo & Son Hy (2024) through Generative AI. These interactions are pivotal for uncovering potential drug candidates, elucidating disease pathways, and crafting effective treatments. Yet, the intricate and diverse nature of molecular biology presents substantial challenges in accurately predicting these interactions, necessitating the development of advanced computational strategies. Recent advancements in deep learning have revolutionized computational biology by offering powerful methods for modeling complex biological systems. Transformer architectures Vaswani et al. (2017), originally designed for natural language processing (NLP), have demonstrated remarkable success in capturing long-range dependencies and intricate relationships. Simultaneously, Large Language Models (LLMs) have shown their ability to generate meaningful embeddings for biological sequences, such as SMILES for drugs Elnaggar et al. (2021); Brandes et al. (2022); Hayes et al. (2025) and amino acid sequences for proteins Chithrananda et al. (2020); Ross et al. (2022); Edwards et al. (2022); Nguyen & Hy (2024); Khang Ngo & Son Hy (2024). These embeddings retain rich biochemical and structural information, providing a promising avenue for understanding molecular interactions.

Existing approaches for molecular interaction prediction leverage various machine learning techniques but face notable limitations. ConPLex Singh et al. (2023) utilizes a pretrained protein language model to predict drug-target interactions by co-locating proteins and drug molecules in a shared feature space, achieving solid performance. However, its reliance on choosing appropriate loss functions based on molecule and protein diversity can lead to data leakage and poor generalization. iNGNN-DTI Sun et al. (2024) applies a nested graph neural network (GNN) for DTI prediction, leveraging pre-trained molecular and protein models, and constructing target graphs from AlphaFold2 3D structures. While it enhances interpretability, it still faces challenges in fully capturing molecular relationships. Its use of a cross-attention-free transformer fails to jointly model drug and protein features due to their distinct distributions, and the reliance on pure MLPs for prediction limits performance, especially in complex cases. SkipGNN Huang et al. (2020) introduces a novel GNN architecture that propagates neural messages via both direct and second-order interactions, improving molecular interaction discovery. However, it still suffers from a lack of biological context, as it does not incorporate pretrained biological language models. MUSE Rao et al. (2024) proposes a multi-scale EM-based framework that integrates structural and network-level information for protein-drug interactions. However, it heavily relies on structural data, which is often unavailable. Moreover, protein structures are merely approximations influenced by experimental techniques like X-ray crystallography Smyth & Martin (2000), cryo-electron microscopy (cryo-EM) Tye et al. (2017), or nuclear magnetic resonance (NMR), all of which introduce uncertainties. In drug design, accurate 3D target structures, such as binding sites, are often missing, further limiting structure-based approaches. Many existing methods, including those by Jha et al. (2022), Zhang et al. (2024), Zhu et al. (2024), and Li et al. (2022), either depend on inaccessible structural data or fail to capture complex relationships due to simplified architectures like MLPs.

To address these challenges, we propose a novel framework that combines the strengths of LLM-based sequence embeddings and Transformer architectures to model molecular interactions. Drugs and proteins are represented as embeddings learned by domain-specific LLMs, capturing their intrinsic biochemical properties. These embeddings are then fused through a Transformer model, which effectively captures the interaction patterns between different molecular entities. This unified approach enables the prediction of DTIs, PPIs, and DDIs with high accuracy and generalization. We evaluate our framework on a diverse set of molecular interaction benchmarks, achieving state-of-the-art (SOTA) performance on multiple DTI and DDI datasets and competitive results on PPI benchmarks. These results highlight the versatility and effectiveness of our approach in addressing various molecular interaction prediction tasks. Our contributions can be summarized as follows:

### Integration of pretrained LLM embeddings with Transformer-based interaction modeling

We combine the rich sequence-level representations of drugs and proteins from pretrained large language models (LLMs) with a Transformer encoder layer to jointly model interactions. This approach captures complex relationships between molecular entities and significantly enhances prediction tasks.

### Broad applicability and SOTA performance

Our method achieves SOTA performance on three standard drug-target interaction (DTI) datasets and competitive results on protein-protein interaction (PPI) and drug-drug interaction (DDI) benchmarks, demonstrating its generalizability and versatility across fundamental biological prediction tasks and making it an ideal candidate for developing foundation models for molecular interaction.

### Efficiency and independence from 3D structural data

Unlike approaches reliant on 3D molecular structures, which are label-intensive and architecturally complex, our framework operates efficiently using only sequence data, making it adaptable to arbitrary drugs and proteins. Its efficacy and versatility make it well-suited for building robust, large-scale foundation models for molecular interactions.

## 2 Method

The Drug-Target Interaction (DTI) prediction task can be framed as a binary classification problem. Given a drug *d* ∈ 𝒟, represented by its SMILES notation, and a protein *p* ∈ 𝒫, represented by its amino acid sequence, the goal is to learn a function *f* : 𝒟 *×* 𝒫 → {0, 1} that predicts whether an interaction exists. Formally, the model aims to approximate *ŷ* = *f* (*d, p*; *θ*), where *ŷ* ∈ {0, 1} represents the predicted interaction (1 for interaction, 0 for no interaction), and *θ* refers to the model’s learnable parameters.

### Drug Representation

The drug *d* is first encoded using a pretrained Small Molecule LLM, denoted as LLM_drug_, which transforms the SMILES notation into an embedding: *h*_*d*_ = LLM_drug_(*d*). This embedding *h*_*d*_ captures the drug’s structural and chemical features. The embedding is then further processed by a Multi-Layer Perceptron (MLP), parameterized by *ϕ*_*d*_, to refine the drug representation: 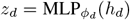. This step enhances the drug feature vector *z*_*d*_, making it suitable for interaction prediction.

### Protein Representation

Similarly, the protein sequence *p* is encoded by a pretrained Protein LLM, denoted as LLM_protein_, which generates a sequence-level embedding: *h*_*p*_ = LLM_protein_(*p*). The embedding *h*_*p*_ represents the protein’s sequence and structure. This sequence-level embedding is further refined by another MLP, parameterized by 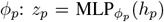. The result is a feature vector *z*_*p*_ that encapsulates the protein’s characteristics relevant to binding with drugs.

### Unified Representation

The drug and protein embeddings are then concatenated to form a unified feature representation: *z*_fusion_ = *z*_*d*_ ⊕ *z*_*p*_, where ⊕ denotes the concatenation operation, combining the drug and protein features into a single vector. The fused representation *z*_fusion_ is then processed by a Transformer encoder 𝒯, which captures complex relationships between the drug and protein features. The attention mechanism in the Transformer allows the model to learn dependencies between the two types of entities (drug and protein), improving the accuracy of predictions: *z*_trans_ = 𝒯 (*z*_fusion_). Finally, a classifier 𝒞, typically implemented as an MLP followed by a sigmoid activation function, is applied to the Transformer output to predict the probability of interaction: *ŷ* = *σ*(𝒞 (*z*_trans_)), where *σ*(·) is the sigmoid function, ensuring that the output *ŷ* is in the range (0, 1), representing the probability of interaction.

### Optimization

The model is trained using binary cross-entropy loss, which measures the difference between the predicted interaction probabilities and the true labels *y*_*i*_ for the *i*-th drug-target pair. The loss function is:

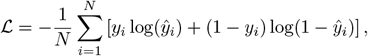

where *N* is the total number of training samples, *y*_*i*_ is the ground truth label for the *i*-th pair (1 for interaction, 0 for no interaction), and *ŷ*_*i*_ is the predicted probability for the interaction. This loss function encourages the model to make accurate predictions by penalizing errors in both positive and negative classifications.

### 2.1 Generalization to Other Interaction Tasks

The proposed architecture is not limited to drug-target interaction prediction. It can be generalized to other interaction prediction tasks, such as protein-protein and drug-drug interactions. For protein-protein interactions, both the Small Molecule LLM and the Protein LLM in Figure 1 are replaced with two instances of the Protein LLM, enabling the extraction of biologically meaningful features from both input protein sequences. Similarly, for drug-drug interactions, the architecture employs two Small Molecule LLMs to process the SMILES representations of the interacting drugs.

**Figure 1:**
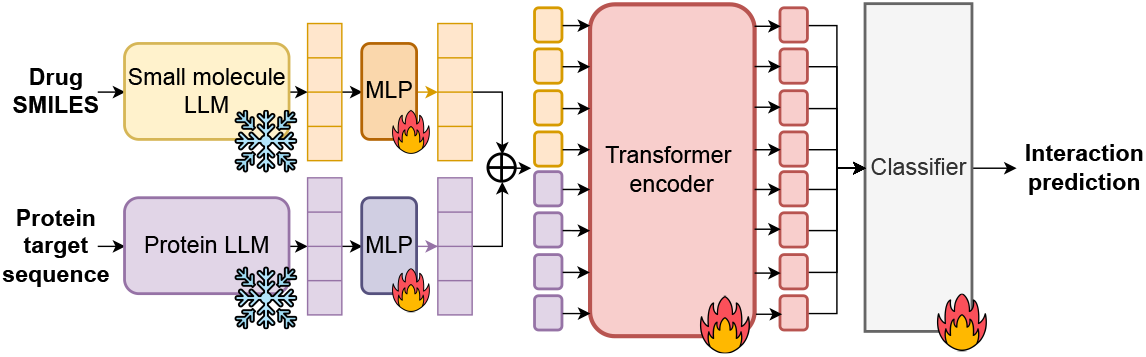
The diagram illustrates the flow of data through different components for the Drug-Target Interaction (DTI) task: Drug SMILES are processed by a Small molecule LLM and then passed through an MLP (Multi-Layer Perceptron). Similarly, Protein target sequences are processed by a Protein LLM and then passed through another MLP. The outputs of these MLPs are concatenated (denoted by ⊕) and fed into a Transformer encoder, which then passes the processed data to a Classifier for interaction prediction.

## 3 Experiments

### 3.1 Datasets

The study leverages a variety of benchmark datasets to assess the performance of the proposed methods in different interaction prediction tasks. For Drug-Target Interaction (DTI) prediction, the DAVIS Davis et al. (2011), KIBA He et al. (2017), and BioSNAP Zitnik et al. (2018) datasets provide comprehensive drug-protein interaction data, summarized in Table 1. For Protein-Protein Interaction (PPI) tasks, the Yeast PPI Ito et al. (2001) dataset is employed, containing 2,497 proteins and 11,188 interactions. In the case of Drug-Drug Interaction (DDI) prediction, the DeepDDI Rao et al. (2024) dataset provides data on drug-drug interactions and side effects, sourced from DrugBank Wishart et al. (2018), making it an essential resource for studying potential adverse drug reactions.

**Table 1:**
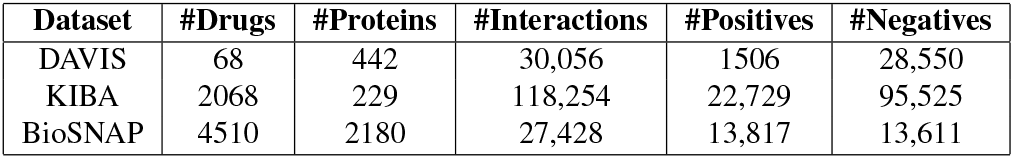
Summary of the DAVIS, KIBA, and BioSNAP datasets used for drug-target interaction (DTI) tasks, including the number of drugs, proteins, interactions, and the distribution of positive and negative interactions.

| Dataset | #Drugs | #Proteins | #Interactions | #Positives | #Negatives |
| --- | --- | --- | --- | --- | --- |
| DAVIS | 68 | 442 | 30,056 | 1506 | 28,550 |
| KIBA | 2068 | 229 | 118,254 | 22,729 | 95,525 |
| BioSNAP | 4510 | 2180 | 27,428 | 13,817 | 13,611 |

### 3.2 Architecture and Implementation Details

The detailed architectural configurations, hyperparameter selections, and implementation specifics are thoroughly documented in Appendix A.1. This section covers critical aspects such as the computational resources utilized, including the type of GPUs employed, the selection of large language models (LLMs), optimization strategies, and the number of layers in the proposed architecture.

### 3.3 Ablation study

BioSNAP and DAVIS were selected for the ablation study due to their distinct characteristics and computational feasibility. DAVIS, which consists of a small number of drugs with dense interaction data, provides a controlled setting for evaluating model performance on well-characterized targets. In contrast, BioSNAP offers a more balanced distribution of positive and negative interactions, enabling a comprehensive assessment of model generalization.

#### Effect of LLMs selection

To identify the optimal large language model (LLM) for interaction prediction tasks, we conducted an ablation study evaluating the performance of various LLMs tailored for proteins and drugs. For proteins, we considered models such as ProtT5 Elnaggar et al. (2021), ProtBERT Brandes et al. (2022) and ESM3 Hayes et al. (2025) (Evolutionary Scale Modeling version 3), while for drugs, we evaluated models like ChemBERTa Chithrananda et al. (2020), MoLFormer Ross et al. (2022) and MolT5 Edwards et al. (2022). The evaluation was performed on two benchmark datasets, DAVIS and BioSNAP, which provide diverse and complementary data for drug-target interaction prediction tasks. The results from these datasets allowed for a robust comparison of the LLMs, highlighting their strengths and weaknesses in capturing biochemical interactions effectively.

Table 2 illustrates the performance of various large language models (LLMs) for proteins and drugs was evaluated on the BioSNAP and DAVIS datasets using AUROC and AUPRC metrics. The results indicate that the combination of ProtT5 and MoLFormer achieved the highest performance on BioSNAP, with AUROC and AUPRC values of 0.9953 and 0.9961, demonstrating its effectiveness in predicting interactions in this dataset. On the DAVIS dataset, ESM3 paired with MoLFormer emerged as the top-performing model, attaining the highest AUROC (0.995) and AUPRC (0.905). MoLFormer consistently delivered strong results across both datasets, showcasing its robustness as a drug representation model. Among the protein models, ProtT5 and ESM3 were particularly effective, with ESM3 excelling in DAVIS and ProtT5 in BioSNAP. Additionally, ProtBERT paired with MoLFormer also showed competitive performance, making it a viable alternative for specific scenarios. Based on these findings, ProtT5 with MoLFormer is recommended for BioSNAP, while ESM3 with MoLFormer is best suited for DAVIS. For applications requiring a single versatile combination, MoLFormer paired with either ProtT5 or ESM3 provides a robust solution.

**Table 2:** Performance comparison of large language models (LLMs) for proteins and drugs on the BioSNAP and DAVIS datasets. The metrics reported are the Area Under the ROC Curve (AU-ROC) and the Area Under the Precision-Recall Curve (AUPRC). The highest performance values are marked in bold, the second-highest are <u>underlined</u>, and the third-highest are *italicized*, showcasing the relative effectiveness of each model combination.

| Proteins | Drugs | BioSNAP |  | DAVIS |  |
| --- | --- | --- | --- | --- | --- |
| | | AUROC (↑) | AUPRC (↑) | AUROC (↑) | AUPRC (↑) |
| ProtT5 | ChemBERTa | 0.9911 | 0.9907 | 0.989 | 0.827 |
|  | MolFormer | **0.9953** | **0.9961** | *0.991* | *0.878* |
|  | MolT5 | 0.9946 | 0.9955 | 0.991 | 0.872 |
| ProtBERT | ChemBERTa | 0.9807 | 0.9857 | 0.990 | 0.849 |
|  | MolFormer | *0.9947* | *0.9953* | 0.991 | 0.857 |
|  | MolT5 | 0.9842 | 0.9880 | 0.991 | 0.867 |
| ESM3 | ChemBERTa | 0.9927 | 0.9938 | 0.990 | 0.871 |
|  | MolFormer | <u>0.9948</u> | <u>0.9958</u> | **0.995** | **0.905** |
|  | MolT5 | 0.9887 | 0.9908 | <u>0.993</u> | <u>0.882</u> |

#### Impact of Transformer-based Encoding

We evaluated the performance of Transformer architectures against traditional Multi-Layer Perceptrons (MLPs) for encoding tasks. Our findings indicate that transformer-based models with the combination of ProtT5 for protein encoding and MoL-Former for drug encoding, significantly outperformed MLP-based approaches on the BioSNAP dataset. Table 3 presents the performance metrics of different model configurations, highlighting the effect of removing or replacing Transformer components. The results demonstrate that the full Transformer-based model achieves the highest AUROC (0.9953) and AUPRC (0.9961), along with superior sensitivity (0.9511) and specificity (0.9562).

**Table 3:**
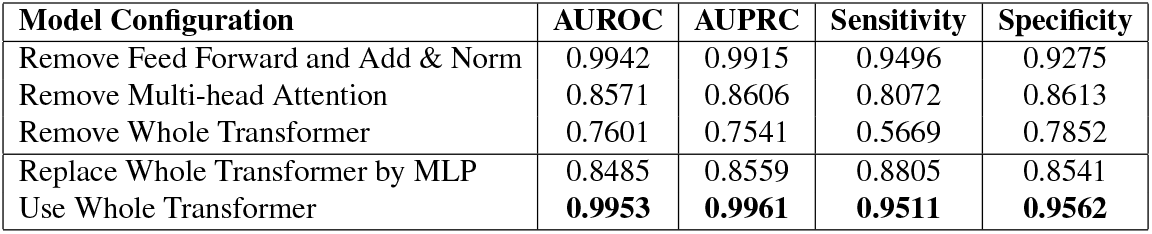
Performance metrics (AUROC and AUPRC) for different model configurations, evaluating the impact of removing or replacing Transformer components.

| Model Configuration | AUROC | AUPRC | Sensitivity | Specificity |
| --- | --- | --- | --- | --- |
| Remove Feed Forward and Add & Norm | 0.9942 | 0.9915 | 0.9496 | 0.9275 |
| Remove Multi-head Attention | 0.8571 | 0.8606 | 0.8072 | 0.8613 |
| Remove Whole Transformer | 0.7601 | 0.7541 | 0.5669 | 0.7852 |
| Replace Whole Transformer by MLP | 0.8485 | 0.8559 | 0.8805 | 0.8541 |
| Use Whole Transformer | **0.9953** | **0.9961** | **0.9511** | **0.9562** |

Ablation studies reveal a substantial decline in performance when key Transformer components are removed. Eliminating the Feed Forward and the Add & Normalization steps results in a slight reduction in AUROC (0.9942) and AUPRC (0.9915), suggesting that these components contribute to fine-tuning the model’s performance. In contrast, removing the Multi-head Attention mechanism causes a significant drop in AUROC (0.8571) and AUPRC (0.8606), emphasizing the crucial role of attention mechanism in feature extraction. The most severe degradation is observed when the entire

Transformer structure is removed, leading to an AUROC of 0.7601 and an AUPRC of 0.7541, with markedly reduced sensitivity (0.5669).

Furthermore, replacing the entire Transformer with a MLP results in an AUROC of 0.8485 and an AUPRC of 0.8559, which is still lower than any Transformer-based configuration, reinforcing the superiority of Transformer architectures over traditional MLPs for encoding tasks. These findings underscore the critical role of Transformer components, particularly Multi-head Attention, in achieving optimal performance in drug-protein interaction prediction.

Appendix A.2 provides an in-depth analysis of the advantages of Transformer-based fusion models compared to traditional MLPs. The results presented in Table 3 further corroborate these findings, emphasizing the critical role of various Transformer components in encoding performance. Specifically, the removal of key elements, such as Multi-head Attention, led to a substantial decline in predictive accuracy, thereby reinforcing the theoretical insights discussed in Appendix A.2.

Figure 2 presents t-SNE visualizations of data representations, illustrating the impact of MLP and Transformer models on feature distribution and clustering. Observing the transformations, it is evident that the MLP modifies the data distribution to a certain extent, enhancing separation but still exhibiting some overlap. In contrast, the Transformer model demonstrates a more pronounced clustering effect, indicating its superior capability in capturing complex relationships and structural patterns within the data. These visualizations underscore the effectiveness of Transformer-based models in producing well-defined feature representations compared to traditional MLP approaches.

**Figure 2:**
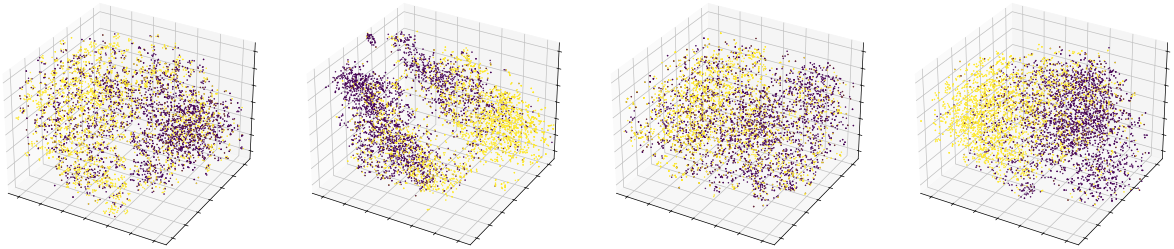
t-SNE visualizations of data representations: (a) before applying the MLP, (b) after applying the MLP, (c) before applying the Transformer, and (d) after applying the Transformer. Data points are color-coded according to their labels, where 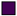 represents label 0, 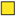 represents label 1.

By integrating both theoretical analysis and empirical validation, our study underscores the enhanced capability of Transformer-based models in capturing complex feature interactions, presenting a compelling case for their adoption in drug-target interaction tasks.

### 3.4 Benchmarks

#### 3.4.1 Drug-target interaction benchmark

Tables 4, 5, and 6 present a comprehensive performance comparison of various drug-target interaction (DTI) prediction methods across the BioSNAP, DAVIS, and KIBA datasets. The results highlight the superiority of our Transformer-based approach over existing models, including DeepDTA Ö ztürk et al. (2018), Moltrans Huang et al. (2021), ML-DTI Yang et al. (2021), DGraphGTA Jiang et al. (2020), and iNGNN-DTI Sun et al. (2024).

**Table 4:** Performance comparison of various methods on the DTI task using the BioSNAP datasets. The table reports the AUROC and AUPRC with their respective standard deviations.

| Method | AUROC | AUPRC | Sensitivity | Specificity |
| --- | --- | --- | --- | --- |
| DeepDTA | $0.897 \pm 0.0027$ | $0.900 \pm 0.0046$ | $0.859 \pm 0.0089$ | $0.786 \pm 0.0197$ |
| Moltrans | $0.887 \pm 0.0034$ | $0.881 \pm 0.0085$ | $0.824 \pm 0.0106$ | $0.809 \pm 0.0104$ |
| ML-DTI | $0.911 \pm 0.0053$ | $0.911 \pm 0.0112$ | $0.851 \pm 0.0054$ | $0.828 \pm 0.0215$ |
| DGraphGTA (Alphafold2) | $0.913 \pm 0.0022$ | $0.917 \pm 0.0024$ | $0.858 \pm 0.0175$ | $0.831 \pm 0.0151$ |
| iNGNN-DTI | $0.934 \pm 0.0021$ | $0.939 \pm 0.0022$ | $0.872 \pm 0.0189$ | $0.854 \pm 0.0200$ |
| Our method | **$0.995 \pm 0.0045$** | **$0.996 \pm 0.0036$** | **$0.951 \pm 0.0409$** | **$0.956 \pm 0.0329$** |

**Table 5:** Performance comparison of various methods on the DTI task using the DAVIS datasets. The table reports the AUROC and AUPRC with their respective standard deviations.

| Method | AUROC | AUPRC | Sensitivity | Specificity |
| --- | --- | --- | --- | --- |
| DeepDTA | $0.892 \pm 0.0066$ | $0.378 \pm 0.0231$ | $0.854 \pm 0.0066$ | $0.792 \pm 0.0291$ |
| Moltrans | $0.898 \pm 0.0050$ | $0.371 \pm 0.0067$ | $0.865 \pm 0.0050$ | $0.783 \pm 0.0387$ |
| ML-DTI | $0.910 \pm 0.0034$ | $0.381 \pm 0.0247$ | $0.895 \pm 0.0034$ | $0.795 \pm 0.0183$ |
| DGraphGTA (Alphafold2) | $0.885 \pm 0.0099$ | $0.316 \pm 0.0447$ | $0.894 \pm 0.0034$ | $0.724 \pm 0.0467$ |
| iNGNN-DTI | $0.931 \pm 0.0027$ | $0.473 \pm 0.0167$ | $0.922 \pm 0.0155$ | $0.802 \pm 0.0240$ |
| Our method | **$0.995 \pm 0.0037$** | **$0.905 \pm 0.0238$** | **$0.976 \pm 0.0159$** | **$0.964 \pm 0.0207$** |

**Table 6:**
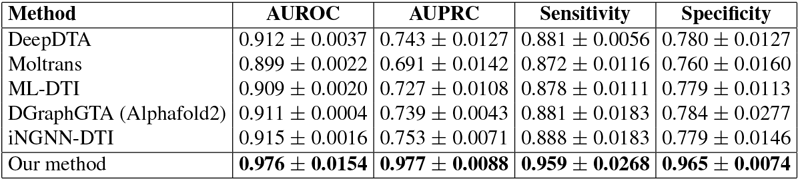
Performance comparison of various methods on the DTI task using the KIBA datasets. The table reports the AUROC and AUPRC with their respective standard deviations.

Across all datasets, our method consistently achieves the highest AUROC and AUPRC, with substantial improvements over state-of-the-art approaches. Notably, on the BioSNAP dataset, our model attains an AUROC of 0.995 and an AUPRC of 0.996, surpassing iNGNN-DTI, the second-best model, by a significant margin. Similar trends are observed for the DAVIS dataset, where our method outperforms competing models with a notable increase in both AUROC (0.995) and AUPRC (0.905). The most pronounced performance gain is seen on the KIBA dataset, where our approach achieves an AUROC of 0.976 and an AUPRC of 0.977, demonstrating its robustness across different benchmarks.

These results underscore the effectiveness of our model in accurately capturing drug-protein interactions, significantly outperforming MLP-based and graph-based methods. The observed improvements in sensitivity and specificity further validate the model’s reliability in both detecting positive interactions and minimizing false positives, making it a promising tool for DTI prediction.

#### 3.4.2 Drug-drug interaction benchmark

Table 7 presents a comparative analysis of drug-drug interaction (DDI) prediction methods on the DeepDDI dataset, evaluated using AUROC and AUPRC metrics. Our proposed method achieves an AUROC of 0.998 and an AUPRC of 0.995, outperforming all other approaches. Notably, while MUSE attains the same AUROC, our method achieves the highest AUPRC, demonstrating superior precision in ranking positive interactions. These results highlight the effectiveness of our approach in enhancing DDI prediction accuracy.

**Table 7:**
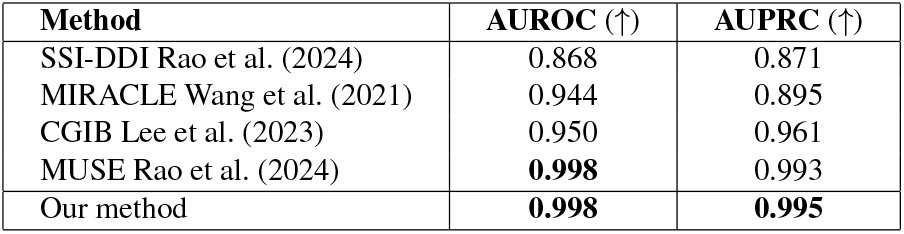
Performance comparison of drug-drug interaction prediction methods using the DeepDDI datasets, evaluated by AUROC and AUPRC metrics. Bold values indicate the best performance in each category.

#### 3.4.3 Protein-protein interaction benchmark

Table 8 presents a comparative evaluation of various methods on the Yeast PPI dataset across multiple performance metrics, including accuracy (Acc), precision (Pre), sensitivity (Sen), specificity (Spe), F1-score (F1), Matthews correlation coefficient (MCC), and area under the curve (AUC). The TAGPPI method demonstrates superior overall performance, achieving the highest scores in most metrics. In contrast, our proposed methods, particularly *ProtBERT* and *ProtT5*, exhibit exceptional sensitivity, reaching a near-perfect 99.82%, along with competitive AUC values. However, this comes at the cost of lower precision and accuracy, making our approach particularly well-suited for applications that prioritize high recall, such as identifying novel protein-protein interactions.

**Table 8:**
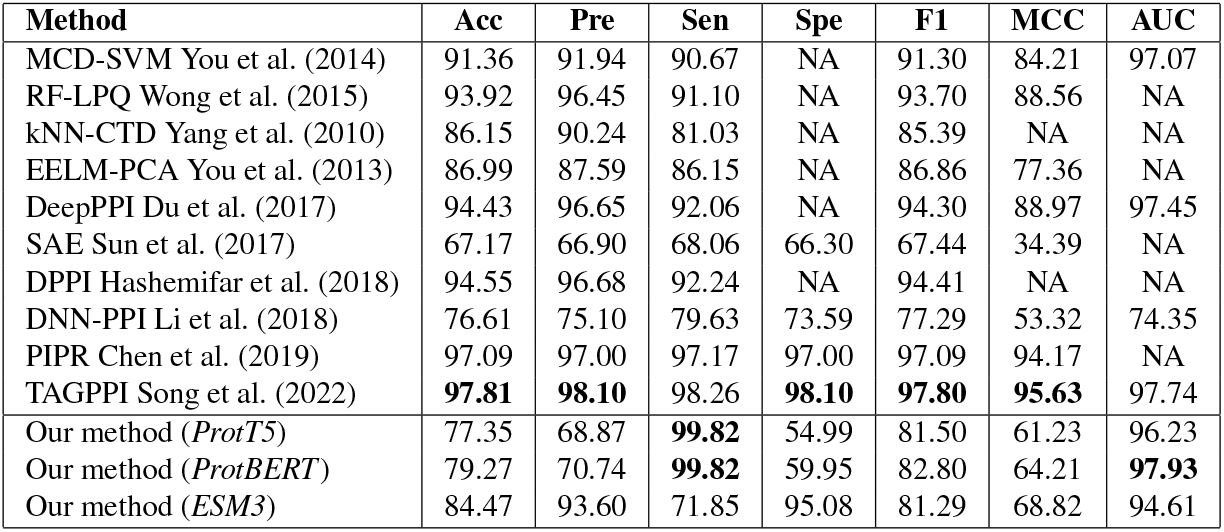
Performance comparison of different methods using the Yeast PPI datasets. NA means the corresponding metric is not available from the original paper. Bold font indicates the best result in the column.

| Method | Acc | Pre | Sen | Spe | F1 | MCC | AUC |
| --- | --- | --- | --- | --- | --- | --- | --- |
| MCD-SVM You et al. (2014) | 91.36 | 91.94 | 90.67 | NA | 91.30 | 84.21 | 97.07 |
| RF-LPQ Wong et al. (2015) | 93.92 | 96.45 | 91.10 | NA | 93.70 | 88.56 | NA |
| kNN-CTD Yang et al. (2010) | 86.15 | 90.24 | 81.03 | NA | 85.39 | NA | NA |
| EELM-PCA You et al. (2013) | 86.99 | 87.59 | 86.15 | NA | 86.86 | 77.36 | NA |
| DeepPPI Du et al. (2017) | 94.43 | 96.65 | 92.06 | NA | 94.30 | 88.97 | 97.45 |
| SAE Sun et al. (2017) | 67.17 | 66.90 | 68.06 | 66.30 | 67.44 | 34.39 | NA |
| DPPI Hashemifar et al. (2018) | 94.55 | 96.68 | 92.24 | NA | 94.41 | NA | NA |
| DNN-PPI Li et al. (2018) | 76.61 | 75.10 | 79.63 | 73.59 | 77.29 | 53.32 | 74.35 |
| PIPR Chen et al. (2019) | 97.09 | 97.00 | 97.17 | 97.00 | 97.09 | 94.17 | NA |
| TAGPPI Song et al. (2022) | **97.81** | **98.10** | 98.26 | **98.10** | **97.80** | **95.63** | 97.74 |
| Our method (*ProtT5*) | 77.35 | 68.87 | **99.82** | 54.99 | 81.50 | 61.23 | 96.23 |
| Our method (*ProtBERT*) | 79.27 | 70.74 | **99.82** | 59.95 | 82.80 | 64.21 | **97.93** |
| Our method (*ESM3*) | 84.47 | 93.60 | 71.85 | 95.08 | 81.29 | 68.82 | 94.61 |

## 4 Conclusion

In this work, we introduced **LANTERN**, a novel framework that leverages Large Language Models (LLMs) and Transformer architectures to enhance molecular interaction prediction. By integrating pretrained embeddings with a Transformer-based fusion mechanism, LANTERN effectively captures complex biochemical relationships across diverse interaction tasks, including Drug-Target Interactions (DTI), Protein-Protein Interactions (PPI), and Drug-Drug Interactions (DDI).

Our extensive evaluations demonstrate that LANTERN achieves state-of-the-art (SOTA) performance on multiple DTI and DDI benchmarks and exhibits competitive results in PPI tasks, underscoring its robustness and versatility. The ablation studies further highlight the importance of Transformer-based encoding over traditional MLP architectures, validating the superior representation learning capabilities of attention mechanisms.

Beyond achieving strong predictive performance, LANTERN offers a scalable and generalizable solution that does not require 3D structural data, making it highly applicable for drug discovery, therapeutic development, and network biology. Integrating self-supervised pretraining strategies may enhance adaptability to new molecular interaction domains. Another key direction is leveraging multiple LLM models within the same data type, such as drugs, to harness the complementary knowledge from various models, addressing individual model limitations and optimizing predictions for a more comprehensive understanding of the problem.

## A Appendix

### A.1 Implementation Details and Model Configurations

In this section, we provide a comprehensive overview of the implementation details and model configurations of the proposed LANTERN framework.

#### A.1.1 LLM choosing

To featurize the inputs, we utilize Molformer, ChemBERTa, and MolT5 for small molecules (embedding sizes per SMILES string: 768, 384, and 1024, respectively) and ProtBERT, ProtT5, and ESM-3 for proteins (embedding sizes per amino acid: 1024, 1024, and 1536, respectively). Our framework is designed to be flexible, allowing for various biological pretrained LLMs, and we propose the choice of LLMs for each task based on empirical observations.

For most language models, we use the output from their final embedding layer. However, for ProtT5 and MolT5, we specifically use the final embedding layer of their encoder components. All models generate per-amino-acid features for proteins or per-SMILES string features for small molecules. These features are averaged along the sequence length to produce fixed-length vectors for downstream tasks.

##### DTI Benchmark

Based on performance across different datasets, we use ProtT5 for proteins in BioSNAP, ProtBERT for proteins in KIBA, and ESM3 for proteins in DAVIS. For small molecules, MoLFormer is selected as the best-performing model across all datasets.

##### DDI Benchmark

MoLFormer is selected due to its robust and consistent performance across experiments.

##### PPI Benchmark

ProtT5, ProtBERT, and ESM3 are all utilized in this task.

##### A.1.2 Evaluation Metrics

Model performance was evaluated using standard metrics, including the accuracy (Acc), precision (Pre), sensitivity (Sen), specificity (Spe), F1-score (F1), Matthews correlation coefficient (MCC), area under the curve (AUC) and area under the precision-recall curve (AUC-PR). The formulas for these metrics are as follows:

**Accuracy (Acc)**measures the proportion of correctly classified samples:

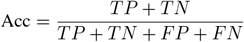

**Precision (Pre)** represents the proportion of true positives among all predicted positives:

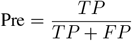

**Sensitivity (Sen)** or **Recall** measures the ability to correctly identify positive samples:

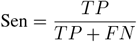

**Specificity (Spe)** measures the ability to correctly identify negative samples:

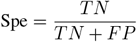

**F1-Score (F1)** is the harmonic mean of precision and sensitivity:

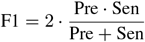

**Matthews Correlation Coefficient (MCC)** is a balanced measure that considers all four quadrants of the confusion matrix:

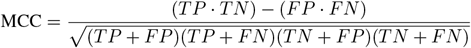

**Area Under the ROC Curve (AUC)** evaluates the trade-off between sensitivity and specificity across thresholds. While no specific formula is shown here, it is calculated using the area under the Receiver Operating Characteristic (ROC) curve.

**Area Under the Precision-Recall Curve (AUC-PR)**: Similar to AUC, but focuses on precision-recall trade-offs, especially useful for imbalanced datasets.

##### A.1.3 Hardware and Software Environment

Experiments were conducted on a system equipped with NVIDIA A100 GPUs and 160GB RAM. The implementation was carried out using PyTorch, with additional libraries such as Hugging Face Transformers for embedding extraction and RDKit for molecular processing.

##### A.1.4 Hyperparameter Tuning

For all three tasks, a linear layer is employed to project the pretrained embeddings from the large language models (LLMs), including the protein language model and the small molecule language model, into a shared latent space of size 384. The two projected representations are then concatenated and passed through an Transformer encoder layer with 8 attention heads. The classifier, a linear layer of shape (768, 1), transforms the encoder output into a logit. The model is trained and optimized for 100 epochs, with a learning rate initialized at 1e-4 and subsequently reduced following a linear annealing schedule after 30, 60, and 80 epochs, using a decay coefficient of 0.8. The batch size is set to 64 for all datasets except the DeepDDI dataset, where it is increased to 512. Dropout is set to 0.1 for all datasets, except for the Yeast dataset, where it is increased to 0.2.

#### A.2 Advantages of Transformer-Based Fusion over MLP

The feature fusion step plays a critical role in learning meaningful interactions between drug and protein representations. While traditional approaches such as Multi-Layer Perceptrons (MLPs) offer simple non-linear transformations, they fall short in capturing complex dependencies between input modalities. In contrast, Transformer-based architectures, driven by the attention mechanism, provide several advantages, including the ability to model long-range dependencies, dynamic weighting of input features, and improved representation learning. This section presents a mathematical formulation to justify the superiority of Transformers over MLPs in the proposed architecture.

##### A.2.1 Limitations of MLP-Based Fusion

An MLP processes concatenated drug and protein feature representations via successive linear transformations followed by non-linear activations. Given the concatenated feature vector *z*_fusion_ ∈ ℝ^*d*^, an MLP of *L* layers with weight matrices *W* ^(*l*)^*l* = 1^*L*^ and biases *b*^(*l*)^*l* = 1^*L*^ produces an output representation:

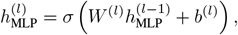

where *σ*(·) denotes a non-linear activation function, such as ReLU. Despite its expressiveness, an MLP applies fixed learned weights to all input features, lacking the flexibility to capture contextual relationships between different segments of the input. Consequently, it struggles with:

- **Static Weighting:** The same weights are applied to all input pairs, disregarding potential interactions between drug and protein features.
- **Lack of Interpretability:** MLPs lack mechanisms to quantify feature importance, making it challenging to understand which features contribute most to the interaction prediction.
- **Poor Scalability:** As input dimensions grow, MLPs require exponentially more parameters to capture feature dependencies effectively.

##### A.2.2 Advantages of Attention Mechanism for Feature Fusion

The self-attention mechanism in Transformers provides a more expressive and efficient method for feature fusion compared to traditional Multi-Layer Perceptrons (MLPs). Unlike MLPs, which apply fixed weights to all input features, self-attention dynamically computes context-dependent feature interactions, leading to improved representation learning. In this section, we provide a mathematical justification of why attention is a more suitable choice for feature fusion in the Drug-Target Interaction (DTI) task.

###### Attention Formulation

Given the concatenated feature representation *z*_fusion_ ∈ ℝ ^*d*^, the selfattention mechanism operates by computing attention scores across all feature dimensions. The attention scores are computed as:

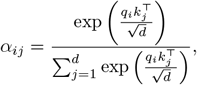

where:

- *q*_*i*_ = *W*_*Q*_*z*_*i*_ and *k*_*j*_ = *W*_*K*_*z*_*j*_ are the query and key projections of the feature representations,
- *W*_*Q*_, *W*_*K*_ ∈ ℝ ^*d×d*^ are learnable weight matrices,
- *α*_*ij*_ represents the attention weight assigned to feature *j* when computing the representation of feature *i*,
- 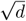 serves as a scaling factor to prevent gradient vanishing.

The output of the attention mechanism is computed as a weighted sum of value vectors:

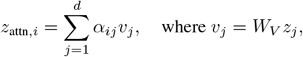

where *W*_*V*_ ∈ ℝ ^*d×d*^ is the learnable value projection.

###### Expressive Power of Attention Compared to MLP

MLPs apply a fixed transformation to the input of the form:

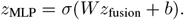

This transformation assumes a linear mapping followed by a non-linear activation *σ*(·), which fails to capture interactions across features in a dynamic fashion. In contrast, attention mechanisms adaptively model feature interactions through:

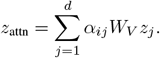

###### Complexity Analysis

Let *d* be the input feature dimension, and assume a hidden dimension of *h*. The computational complexity of an MLP is 𝒪 (*d · h*), whereas the complexity of the self-attention mechanism is 𝒪 (*d*^2^). Although the attention mechanism has a quadratic complexity with respect to input size, the added computational cost is justified by the superior feature interaction modeling capabilities and improved generalization.

Given an arbitrary function *f* : ℝ^*d*^→ ℝ^*d*^, it has been demonstrated that a multihead-attention mechanism, when equipped with a sufficient number of heads and layers, can approximate *f* with lower sample complexity than a Multilayer Perceptron (MLP) of comparable depth. Specifically,for any *ϵ >* 0, the approximation error of attention-based models satisfies the following bound:

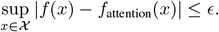

Moreover, the scaling of the approximation error for attention-based models is more favorable than that of MLPs. In particular, the error for MLPs decreases at a rate of 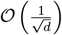, whereas the error in attention-based models diminishes at a faster rate of 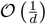

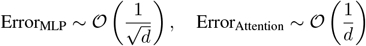

These results indicate that Transformers, which rely on self-attention mechanisms, require fewer parameters to achieve a similar level of approximation accuracy compared to MLPs. This property highlights the parameter efficiency of attention-based models, making them particularly well-suited for high-dimensional input spaces.

###### Empirical Validation

To empirically validate the advantages of attention-based feature fusion, we conducted experiments comparing the Transformer and MLP architectures in terms of model accuracy and loss convergence. The results in Table 3 demonstrate that the attention-based model achieves significantly better performance, supporting the theoretical findings.

###### Conclusion

The self-attention mechanism provides a mathematically superior alternative to MLP-based fusion by offering:

- **Dynamic feature weighting** that adjusts to the importance of different drug and protein features,
- **Efficient long-range dependency modeling** without the need for exponentially large parameters,
- **Stronger generalization abilities** due to better function approximation properties.

Thus, replacing MLPs with attention-based mechanisms in the proposed DTI model enhances feature interaction learning, ultimately leading to improved prediction performance.

Given the above theoretical and empirical justifications, the Transformer-based fusion approach provides a more effective solution for capturing drug-target interactions than MLPs. The proposed model capitalizes on attention mechanisms to dynamically learn intricate feature relationships, leading to improved interaction prediction performance.

